# Investigating the Association Between Nitrate Dosing and Nitrite Generation by the Human Oral Microbiota in Continuous Culture

**DOI:** 10.1101/2023.11.15.567236

**Authors:** Thomas Willmott, Hannah J. Serrage, Elizabeth C. Cottrell, Gavin J. Humphreys, Jenny Myers, Paul M. Campbell, Andrew J. McBain

**Affiliations:** Maternal and Fetal Health Research Centre, Division of Developmental Biology & Medicine, School of Medical Sciences, Faculty of Biology, Medicine and Health, The University of Manchester; Division of Pharmacy and Optometry, School of Health Sciences, Faculty of Biology, Medicine and Health, The University of Manchester

**Keywords:** Oral Microbiota, Multiple Sorbarod Device, Dietary Nitrate, Nitrite, Nitric Oxide, Blood Pressure, Hypertension

## Abstract

The generation of nitrite by the oral microbiota is believed to contribute to healthy cardiovascular function, with oral nitrate reduction to nitrite associated with systemic blood pressure regulation. There is the potential to manipulate the composition or activities of the oral microbiota to a higher nitrate-reducing state through nitrate supplementation. The current study examined microbial community composition and enzymatic responses to nitrate supplementation in in sessile oral microbiota grown in continuous culture. Nitrate reductase activity and nitrite concentrations were not significantly different to tongue-derived inocula in model biofilms. These were generally dominated by *Streptococcus* spp., initially, and a single nitrate supplementation resulted in the increased relative abundance of the nitrate-reducing genera *Veillonella, Neisseria* and *Proteus* spp. Nitrite concentrations increased concomitantly and continued to increase throughout oral microbiota development. Continuous nitrate supplementation, over a 7-day period, was similarly associated with an elevated abundance of nitrate-reducing taxa and increased nitrite concentration in the perfusate. In experiments in which the models were established in continuous low or high nitrate environments, there was an initial elevation in nitrate reductase, and nitrite concentrations reached a relatively constant concentration over time similar to the acute nitrate challenge with a similar expansion of *Veillonella* and *Neisseria*. In summary, we have investigated nitrate metabolism in continuous culture oral biofilms, showing that nitrate addition increases nitrate reductase activity and nitrite concentrations in oral microbiota with the expansion of putatively NaR-producing taxa.

**Importance:** Clinical evidence suggests that blood pressure regulation can be promoted by nitrite generated through the reduction of supplemental dietary nitrate by the oral microbiota. We have utilised oral microbiota models to investigate the mechanisms responsible, demonstrating that nitrate addition increases nitrate reductase activity and nitrite concentrations in oral microbiota with the expansion of nitrate-reducing taxa.

## Introduction

Hypertension, defined as blood pressure (BP) equal to or greater than 140/90 mmHg, significantly increases the risk of haemorrhagic stroke, myocardial infarction, heart failure and chronic renal disease [1, 2]. Between 1995 and 2015, there was a global increase in systolic BP (SBP), loss of disability-adjusted life-years and BP-related deaths [3]. As one of the most preventable causes of premature morbidity and mortality globally, improved treatment and prevention of hypertension is a high priority. Modern genomic-driven diagnostic and therapeutic technologies may offer new biomarkers and novel pathways in the treatment of hypertension [4]. The metabolism of dietary nitrate to nitrite which may regulate systemic cardiovascular function and BP [5-7] offers a novel route for intervention. There is currently substantial ongoing research into using dietary nitrate (as reviewed by Alzahrani, Jackson [8]) to target nitrate reduction to improve systemic cardiovascular health, all reporting wide variability in efficacy between agents and target populations [9, 10].

Inorganic nitrate is introduced into the human body via food and drinking water. Dietary nitrates originate from the fixation of environmental nitrogen by bacteria and are accumulated in high abundance in green, leafy vegetables, such as spinach and lettuce [11]. Additional sources include nitrate/nitrite ions added to cured meats as a preservative [12]. Once ingested, nitrate is rapidly absorbed across the gastrointestinal tract (GI) and into the plasma. Nitrate is actively transported from the plasma into the salivary glands by sialic acid transporters [13, 14], resulting in high nitrate concentrations in saliva, where oral bacteria can reduce the nitrate to nitrite by the actions of nitrate reductase (NaR) enzymes [14]. The surface of the tongue is reported to be the location where most of the nitrate reduction within the mouth occurs [15, 16]. The nitrite then enters the GI tract and leads to the generation of nitric oxide (NO) in blood and tissues, which is dubbed the alternative enterosalivary pathway. NO is a pleiotropic signalling molecule, with wide-ranging effects in the body, and is critical in cardiovascular homeostasis. Reduced bioavailability of NO has been implicated in the pathology of cardiovascular and renal diseases [17-22]. NO regulates vascular tone by relaxing smooth muscle cells, leading to vasodilation of blood vessels. NO therefore has the potential to lower high BP via smooth muscle vasodilation [20].

Disruption of microbial homeostasis in the mouth has been associated with hypertension [23]. It has been reported that disturbances of the oral microbial communities as a result of mouthwash use may decrease oral NaR activity and increase BP in both hypertensive (Bondonno, Liu et al. 2015) and normotensive adults [24-28]. The oral microbiota, therefore, may play a critical role in our systemic health and, consequently, manipulation of these communities may offer a novel tool of intervention in cardiovascular disorders [29]. There is now an emerging field of microbiome-driven research aimed at increasing NO production, via nitrate supplementation, for the treatment of hypertension [29].

There is currently uncertainty about the clinical benefits of dietary supplementation with nitrate. Studies have generally utilised nitrate-rich beetroot juice as the dietary intervention. Some studies reported a significant lowering of BP and an improvement in vascular function in adults [30-34] whilst others reported no effects on cardiovascular function from acute or chronic dietary nitrate dosing [35, 36]. Higher oral NaR is significantly associated with greater BP decrease following acute nitrate supplementation [37-40]. Understanding this variability in the efficacy of nitrate supplementation in the treatment of hypertension may enable the optimisation of interventions. Currently, there is little information about the effects of nitrate supplementation on the taxonomic composition of the oral microbiota. In the present study, therefore, we have used the Multiple Sorbarod Device (MSD) to establish oral microcosms in an *in vitro* continuous culture system and used this to evaluate the effects of nitrate dosing on bacterial community structure and associated NaR activity. MSDs are based on the establishment of biofilms on filters through which growth medium is perfused, which enables both biofilm and perfusate to be monitored. Models were inoculated with samples from the tongue of several healthy volunteers to replicate communities most relevant to the oral microbiome. Bacterial populations were monitored using 16S rRNA gene sequencing. NaR activity and perfusate nitrite concentration were determined longitudinally, following nitrate dosing.

## Materials and Methods

### Volunteers

All reagents were obtained from Sigma-Aldrich (Gillingham, Dorset, UK) unless otherwise stated. Samples were collected following approval from the University of Manchester Research Ethics Committee (Reference: 2019-5929-9344). Healthy participants (n=4; subjects 1-4) were recruited via study advertisements. Inclusion criteria included being aged between 18 and 45, no antibiotic use within 6 months, not currently experiencing any self-assessed oral disease and not a current tobacco smoker.

### Oral sample collection

Tongue surface samples were collected using a sterile Tongue Cleaner (Superdrug, Surrey, UK) in three motions from the rear to the front of the tongue. The end of the Tongue Cleaner was placed in a sterile vessel containing 2mL phosphate-buffered saline (PBS, pH=7.4, M/L= 1.37M sodium chloride, 0.27M potassium chloride, 0.1M phosphate buffer; VWR, Lutterworth, UK) and 1g of 3.5-5.5mm sterile glass beads (VWR, Lutterworth, UK). The solution was mixed via vortex (30 seconds), transferred to a microcentrifuge tube (Thermo Scientific, Massachusetts, USA) and frozen at -80 °C for subsequent analysis.

### Continuous culture oral microcosms

MSDs are *in vitro* models which support the growth of biofilms on five cellulose filter cylinders in parallel (10mm diameter; 20mm length) within a stainless-steel housing. The use of the device to grow oral microcosms has been previously reported [41-45]. MSDs can be sampled destructively by removal and processing of individual colonised filters or by sampling the perfusate which is generated continuously. MSDs were contained within a standard incubator maintained at 36 °C. In this study three models were exploited to 1) validate the use of MSDs for assessment of time-dependent changes in oral microcosm composition and nitrate metabolism (subject 1), 2) explore how nitrate intervention might impact both metabolism and microbial community composition (subject 2) and 3) compare the effects of nitrate intervention relative to a non-treated control over time (subject 3/4). The models utilised are depicted in Figure 1.

**Figure 1:**
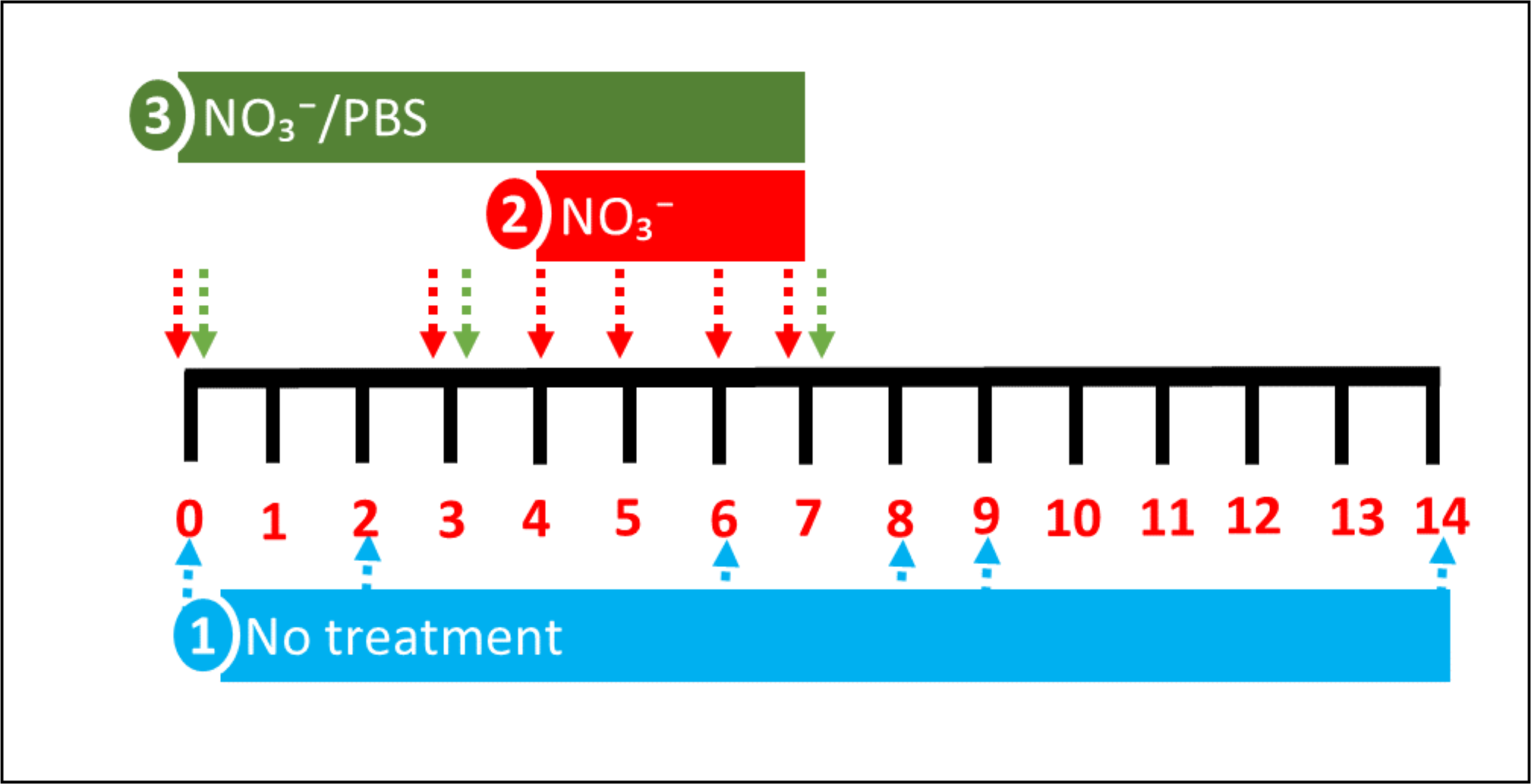
Models utilised to explore nitrate metabolism *in vitro*. Three models were utilised in this study to 1) validate the use of an oral microcosm model to examine time-dependent changes in microbial community composition and nitrate metabolism, 2) explore how nitrate (15mM) intervention manipulates biofilm composition and 3) compare the effects of nitrate (15mM) on biofilm composition and nitrate metabolism relative to a phosphate buffered saline (PBS) control in oral microcosm communities derived from two individuals. Arrows depict time points for sample collection for 16s rRNA sequencing and nitrate metabolism assays.

### Microcosm growth conditions

Microcosms were formed in modified artificial saliva comprising (g/L in distilled water: 2.5g mucin (type II; porcine; gastric), 2.0g bacteriological peptone, 2.0g tryptone, 1.0g yeast extract, 0.35g sodium chloride, 0.2g potassium chloride, 0.2g calcium chloride, 0.1g cysteine hydrochloride, 0.001g haemin, 0.002g vitamin K1) under gentle agitation using a magnetic stirrer [42]. Models were preconditioned with growth medium for 2 h via a peristaltic pump (Minipulse 3; Gilson, Villiers-Le-Bel, France) and throughout model runs (flow rate = 4mL/min.). Pumps were stopped during inoculation and sampling for approximately 10 minutes. Models were inoculated twice, 4 h apart, with 200μL of the oral sample suspension as previously described [46]. Nitrate (15mM; selected based upon the maximum human salivary concentration following dietary nitrate dosing, observed previously; [38]) or phosphate-buffered saline (PBS, control) were applied as depicted in Figure 1 [47]. The inoculum was frozen at -80 °C for subsequent 16S rRNA gene sequencing and nitrate metabolism studies as described below.

**Figure 1:**
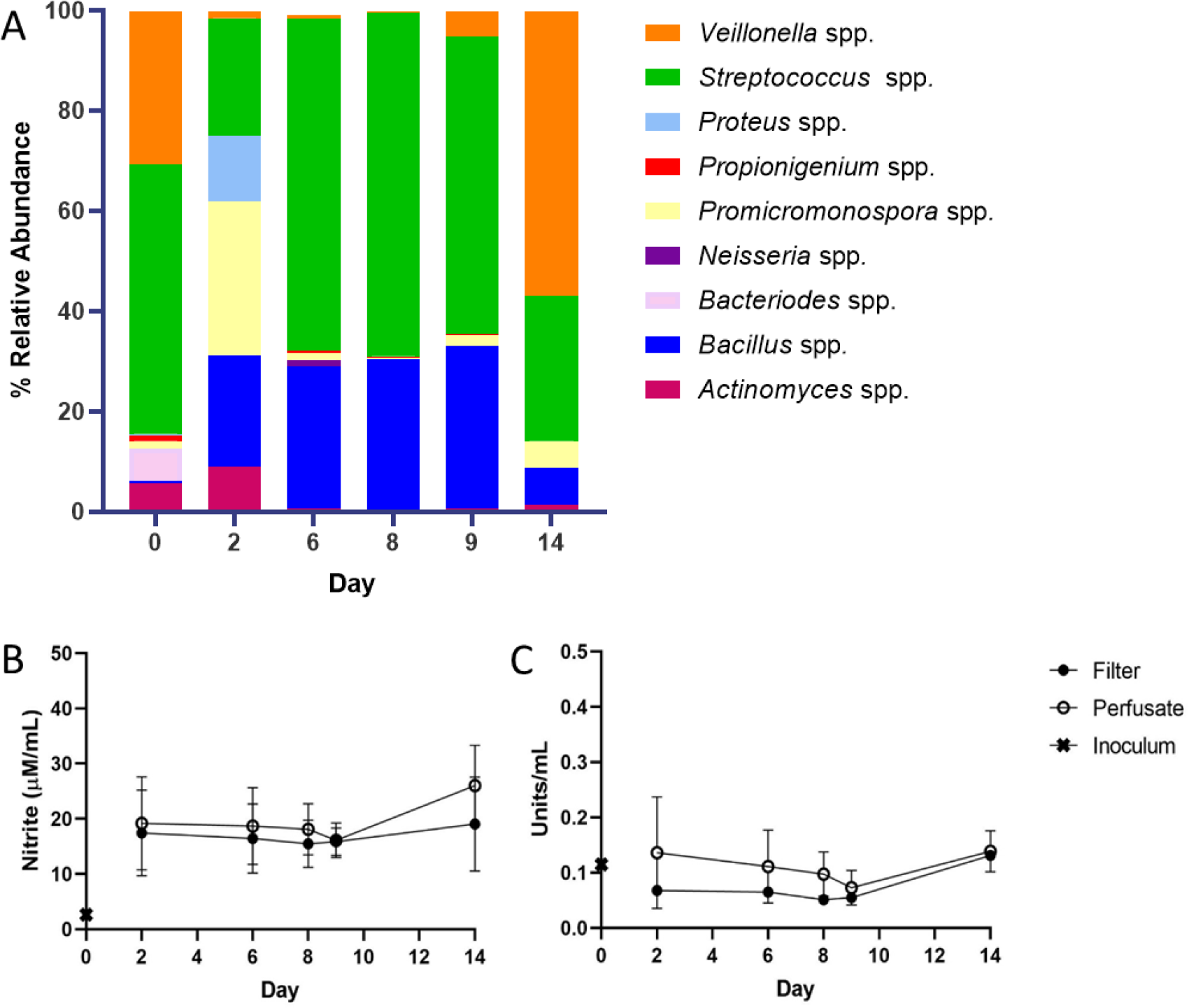
Detectable Levels of Nitrite and Nitrate Reductase in Oral Biofilm Consortia. MSD model was established and inoculated with tongue material collected from subject 1 inoculum. Community composition and nitrate metabolism were assessed via 16S rRNA sequencing (A) and NaR assays (B, C; n = 2) over a 14-day period.

### Analysis of Biofilms and Perfusate

After 3 d of microcosm equilibration, 5mL perfusate (PA) was collected by placing a sterile Universal bottle (Medline Scientific, Oxfordshire, UK) underneath the MSDs waste tube. After sample collection, the area was manually sterilised using ethanol (70% v/v) and the waste collection vessel reattached. To collect the biofilms (BF), the peristaltic pump was halted, and the model was opened aseptically. A single Sorbarod filter was removed and replaced with a sterile filter. Sorbarod filter samples were homogenised via placement in half-strength thioglycolate medium (14.5 g/L; Oxoid, Basingstoke, UK) and 1g of 3.5-5.5mm sterile glass beads (VWR, Lutterworth, UK). Samples were vortexed (30 seconds), transferred to a sterile tube (Thermo Scientific, Massachusetts, USA) and frozen at -80 °C for subsequent molecular analysis.

### Nitrate Reductase (NaR) and Nitrite Assays

The NaR assay was adapted from Lowe and Evans [48] and Allison and Macfarlane [49]. Test inocula (0.1ml) were equilibrated to 36 °C in a buffer comprising 25 mM potassium phosphate buffer, 10 mM potassium nitrate, 0.05 mM ethylenediaminetetraacetic acid (1.8ml) and 2 mM β-nicotinamide adenine dinucleotide (0.1ml; MP Biomedicals, California, USA). In parallel, baseline levels were determined via resuspension of test inocula (0.1ml) in 58 mM (1ml) sulphanilamide stop solution. The baseline and test inocula were then incubated at 36 °C under gentle agitation (100 RPM) for one hour, stop solution was added to the test inocula (2 minutes). Colourimetric assessment was undertaken by the addition of 0.77 mM N-(1-Naphthyl) ethylenediamine dihydrochloride solution to test and baseline samples (10 minutes). Solutions were dispensed into a sterile 96 well plate (Corning, New York, USA), alongside a nitrite standard curve (0-36mM) and absorbance was acquired at 540nm (BioTek, Vermont, USA).

### 16S rRNA gene sequencing and bioinformatics

DNA was extracted from test inocula, Sorbarod filter and MSD perfusate samples using a DNeasy PowerSoil Kit (Qiagen, Hilden, Germany) according to the manufacturer’s instructions. Amplification was achieved through 16S rRNA gene Polymerase Chain Reaction (PCR) with Illumina (San Diego, USA) adapted primers [50] and NEBNext^®^ High-Fidelity 2X PCR Master Mix (New England Biolabs, Ipswich, USA) at 98°C (2 minutes) followed by 25 cycles of 95°C (20 seconds), 62°C (15 seconds), 70°C, (30 seconds) and a final elongation step of 72°C (5 minutes), which have been previously validated in oral microbiome studies [51-54]. The amplified 16S rRNA gene was purified using a QIAquick purification kit (Qiagen, Hilden, Germany) and amplicons confirmed via gel electrophoresis.

Sequencing of the 16S rRNA gene V4 amplicons was performed on the Illumina MiSeq platform (Illumina Inc, Cambridge, UK). Raw sequence data were imported into the quantitative insights into microbial ecology (QIIME) version 2 (2020.2) [55, 56]. Sequences were de-replicated, similarity clustered, analyses for chimaeras, demultiplexed and quality filtered using the d2-demux plug-in followed by denoising with DADA2 (q2-dada2) [57]. Amplicon sequence variants (ASVs) were aligned via the q2-alignment plug-in and taxonomy assigned to the ASVs using the q2-feature-classifier [58] against the Greengenes (v13.8) 97% ASVs reference sequences [59] for the generation of BIOM tables comprising the sample metadata. ASVs present in the negative controls were subtracted from the remaining samples. Following the importation of data using the qiime2R package [60], analysis was performed using the Phyloseq package [61] converting data into relative abundance and ggPlot2 [62] was applied to generate figures in R version 3.6.2 [63]. Statistical investigation of differential analysis of count data was accomplished with DESeq2 [64].

## Results

### Validation of a model for reliable assessment of time-dependent changes in oral microcosm composition *in vitro*

Oral biofilm formation is a dynamic process, in which pioneer colonisers including *Streptococcus* spp. initiate adhesion to the salivary pellicle, facilitating the recruitment of *Veillonella* spp. and subsequent biofilm maturation [65]. We present a model capable of monitoring temporal changes in oral biofilm composition, which indicated diminished diversity relative to the initial inocula (Figure 2A, suppl. Table 1), although this was not significant for either alpha or beta diversity (P>0.05). Microbial community composition was relatively stable across the 14-day culture period, with *Streptococcus* spp. providing most abundant across the study (Figure 2A – C, suppl. Table 1). Over the 14-day period, detectable levels of *Bacillus* spp. were observed and comprised 0.7% of the initial inoculum, the presence of which is associated with yellow tongue coating [66]. Initial study also revealed little temporal change in community composition following 6 -8 days, therefore a 7-day incubation period was selected for nitrate intervention studies.

**Figure 2:**
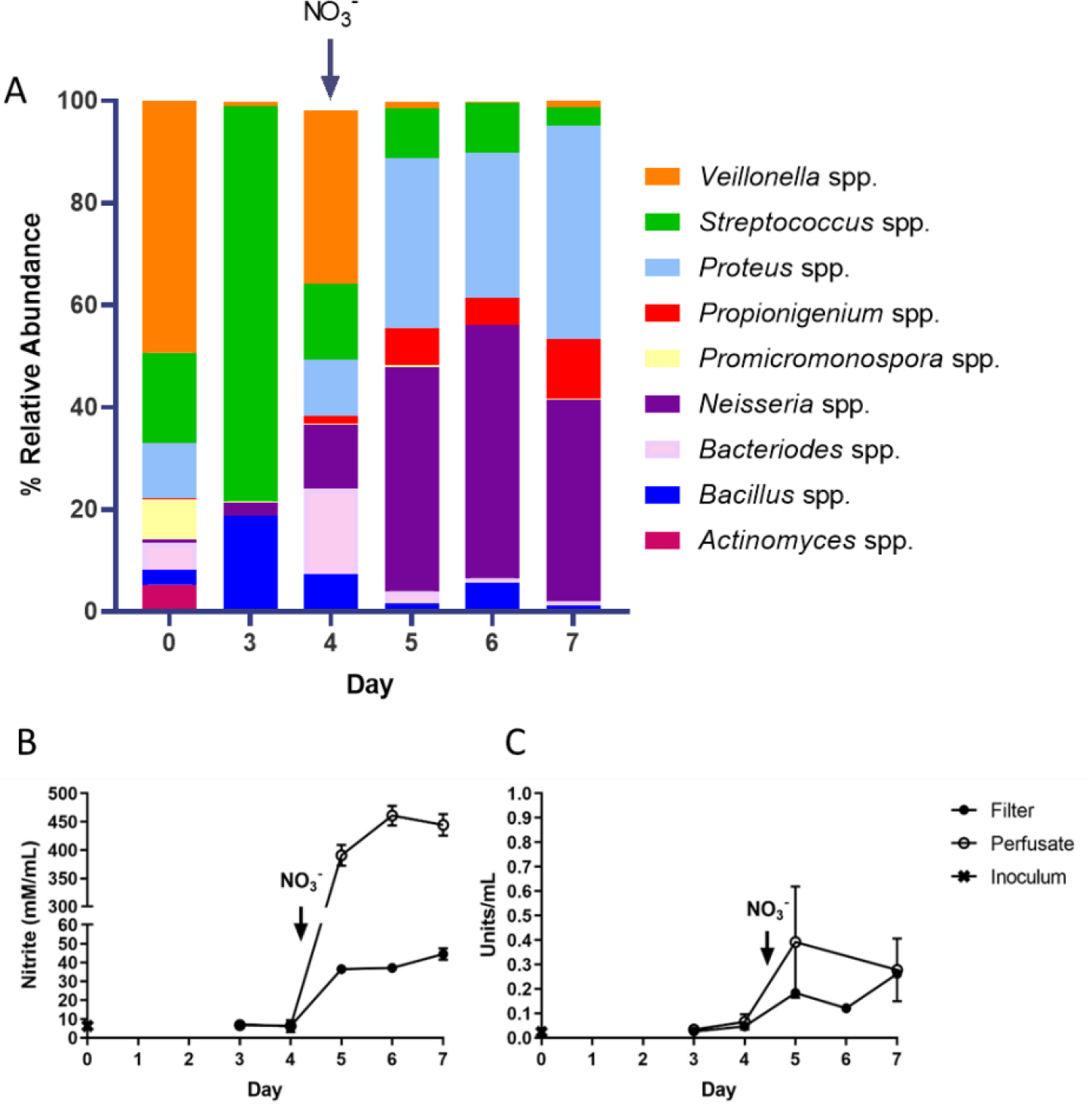
Elevated abundances of *Neisseria* and *Proteus* spp. following exposure of pre-formed biofilms to nitrate. Oral biofilm microcosms derived from subject 2 were established for 4 days on cellulose filters in a continuous culture flow-through system, and subsequently exposed to nitrate. Changes in biofilm composition (A) and nitrate metabolism (B, C) were assessed over a 7-day period from both biofilm (filter) and planktonic (perfusate) samples.

### Exogenous Nitrate Application Elevates Nitrite Production in Filtered but Not Perfused Biofilms

As observed when validating this system, nitrite and nitrate reductase are present at detectable levels in oral microcosm consortia. However, the acute exogenous application of nitrate has proven to not only impact biofilm nitrate metabolism but also composition. Nitrate-reducing genera including *Neisseria* spp. are enriched, whilst periopathogens including *Porphyromonas* spp. are diminished 9h post nitrate application [67, 68]. However, oral biofilm maturation can occur over a period of days [42, 69, 70]. Therefore, nitrate was applied to pre-formed biofilms (4 days) and subsequently, time-dependent changes in nitrate metabolism and biofilm composition were assessed. Intriguingly, nitrate intervention resulted in an increase in nitrite production in collected perfusate (Figure 3B). Nitrate reductase activity also appeared to spike 24h post intervention but stabilised on days 6-7 (Figure 3C). Nitrate application was also accompanied by a shift in community composition (Figure 3A). Prior to nitrate application (day 3), biofilm was dominated by *Streptococcus* spp., but following nitrate intervention on day 4, shifted towards an increasing abundance of *Neisseria* and *Proteus* spp., a finding previously observed by Rosier *et al*. [67]. To verify this shift was due to nitrate intervention, changes in composition and nitrate metabolism were assessed in biofilms treated with nitrate or PBS (control) throughout culture.

**Figure 3:**
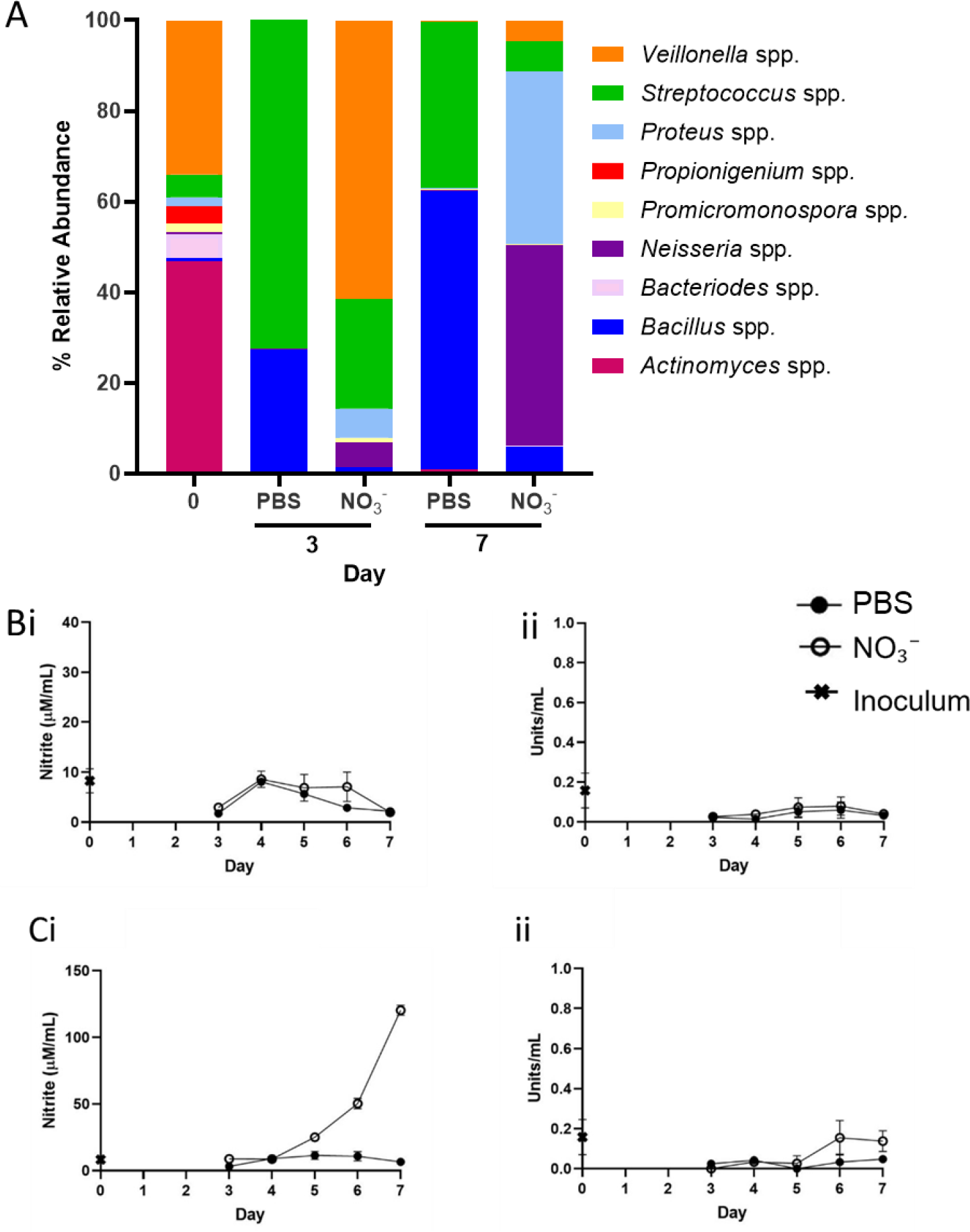
Nitrate Intervention elevates the abundance of *Neisseria* and *Proteus* spp. ASVs. Oral Microcosm (subject 3) were inoculated into the continuous flow-through system and exposed to either 15mM nitrate (NO_3_^-^) or PBS (control) continuously for 7 days. Changes in biofilm composition (A) were assessed via 16s rRNA sequencing. Nitrate metabolism from collected biofilms (filter; B) or planktonic cultures (perfusate; C) were examined via nitrite secretion (i) and nitrate reductase activity (ii) assays.

### Nitrate Intervention May Result in the Elevated Abundance of *Neisseria spp* in Some Individuals

There are significant inter-individual differences in oral microbiota composition, including the abundance of nitrate reducing species such *Neisseria, Rothia* and *Veillonella* [71]. Initial oral microcosm communities in two individuals used in this study proved divergent (Figures 4-5), in which subject 3 (Figure 4A, suppl. Table 3A) exhibited an initial community dominated by *Veillonella* and *Actinomyces* spp. Comparatively, *Actinomyces, Bacteroides, Veillonella* and *Neisseria* spp. proved the most abundant genera of inocula derived from subject 4 (day 0, Figure 5A, suppl. Table 3B). Culture with continuous nitrate appeared to shift the community composition of subject 3, where like in figure 3 (subject 2), on day 7 there was an increased abundance of nitrate-reducing *Proteus* and *Neisseria* spp. relative to the untreated control (Figure 4A). This increased abundance conferred elevated detectable levels of nitrite in perfusate (Figure 4Ci) but not biofilm samples (Figure 4Bi), relative to the PBS control. Comparatively, whilst elevated levels of nitrite secretion could be observed following 5 days of nitrate application in perfusate samples (Figure 5Ci), intervention exerted no significant effect on microcosm composition of subject 4. Whereby *Neisseria* spp. abundance was stable (37.7% ± 5.1) regardless of treatment group on days 3 and 7. Initial inoculum comprised higher levels of *Neisseria* spp. in subject 4 (14.9%) when compared to subject 3 (0.5%). Assessment of pooled ASVs from days 3 and 7 revealed significant increases in *Veillonella* spp. and *Neisseria* spp. following nitrate intervention in subject 3 relative to the PBS control (Figure 6A, p<0.05), but inconsistent effects in subject 4 (Figure 6B). Whereby increases and decreases in *Neisseria* ASVs were observed relative to the PBS control (Figure 6B, p<0.05).

**Figure 4:**
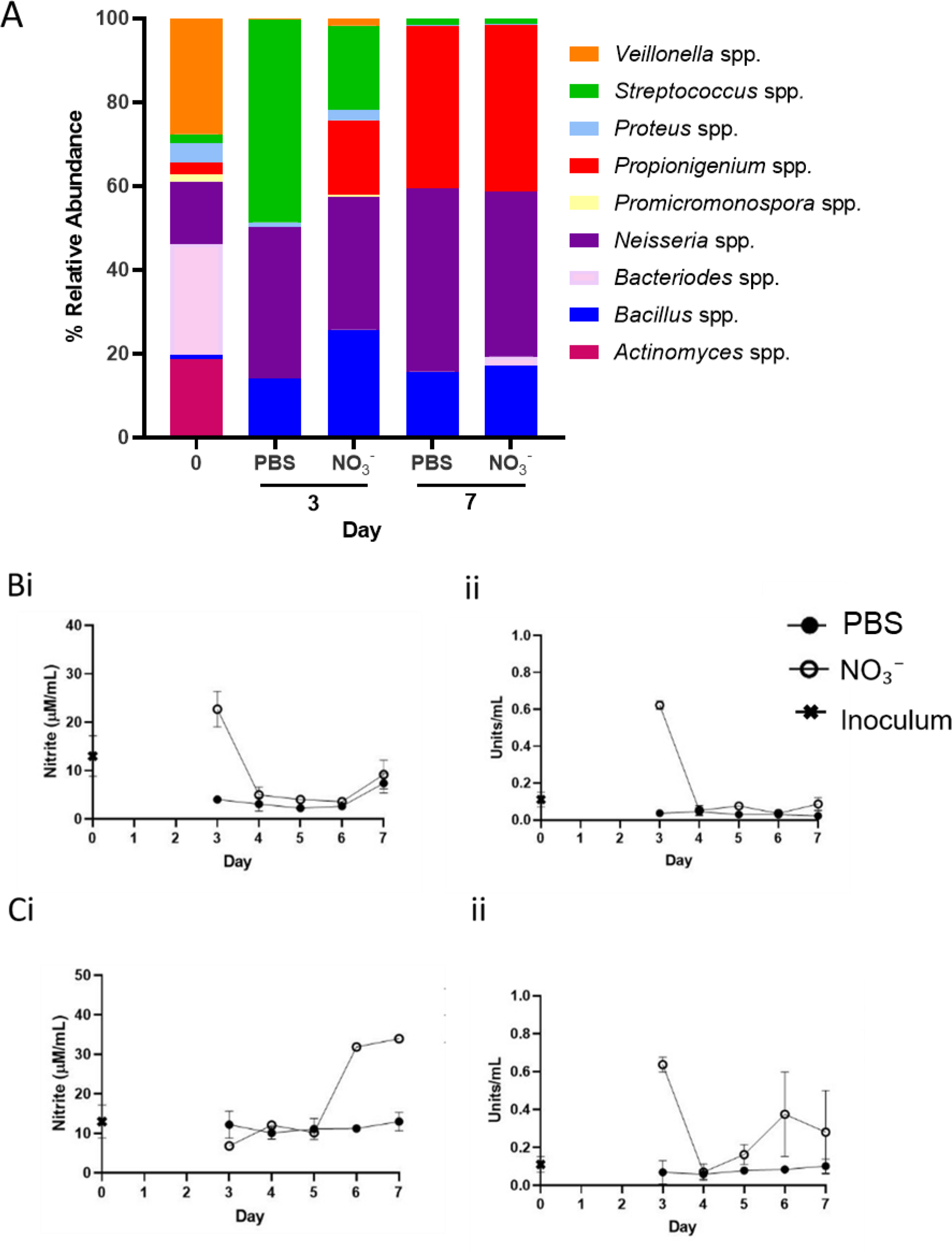
Effects of Nitrate Intervention May Depend Upon Initial Microcosm Composition. An Oral Microcosm (Subject 4) was inoculated into the continuous flow-through system and exposed to either 15mM nitrate (NO_3_^-^) or PBS (Untreated; UT) continuously for 7 days. Changes in biofilm composition (A) were assessed via 16S rRNA sequencing. Nitrate metabolism from collected biofilms (filter; B) or planktonic cultures (perfusate; C) was evaluated via nitrite secretion (i) and nitrate reductase activity (ii) assays.

**Figure 5:**
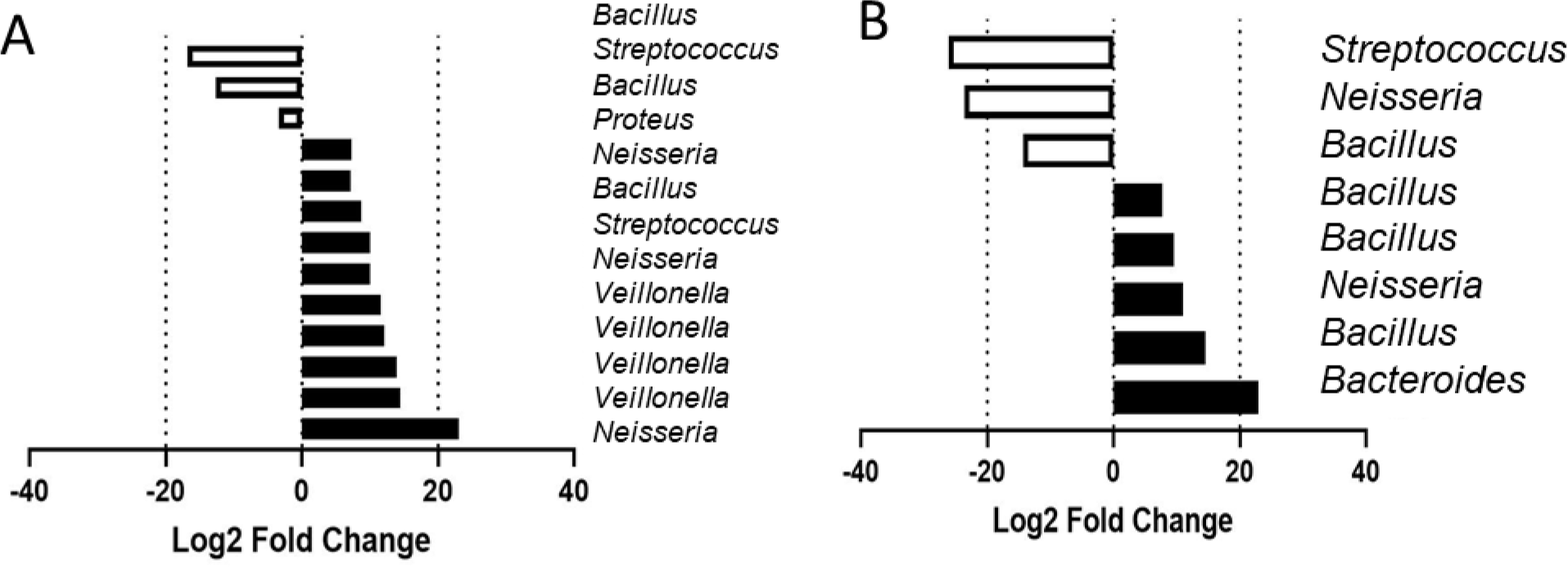
Shift towards high nitrate-reducing organisms following nitrate dosing. Oral Microcosms derived from two individuals (Subject 3; A and Subject 4; B) were either dosed with 15mM nitrate or PBS and data pooled following acquisition of 16s data on collections days 3 and 7. Figures depict combined changes in ASVs on days 3 and 7 in nitrate treated samples, relative to PBS control.

## Discussion

The entero-salivary nitrogen oxide cycle involves the conversion of diet-derived nitrate into nitrite by oral bacteria, and the reduction of nitrite to nitric oxide (NO) in host circulation and tissues [6, 72]. Recent research has identified this metabolic pathway as an important link between oral and systemic health [22, 73]. The composition of the oral microbiota, and the associated nitrate-reducing metabolic potential therefore could provide an opportunity to intervene. We hypothesised that dosing with inorganic nitrate would provide a competitive advantage for certain taxa and may result in their clonal expansion but that responses to nitrate exposure would vary between individual microbiota. Nitrate dosing resulted in a shift in bacterial profiles towards a higher relative abundance of putative nitrate-reducing genera such as *Neisseria*, coupled with an increase in the rate of nitrite formation, that was affected by inter-individual variation in the initial inoculum.

The efficacy of inorganic nitrate supplementation and the downstream effects of the entero-salivary pathway have been reported to vary considerably between individual [73, 74] although the reasons for this variability are not fully understood. In a systematic review of studies investigating the use of inorganic nitrate for treatment of adults with high BP, Remington and Winters [75] reported that of the 12 studies identified, five reported a significant reduction (P<0.05) in both systolic (SBP) and diastolic blood pressure (DBP). One study reported a significant reduction in SBP only, and six reported no reduction in BP. Another review of the role of oral microbiota in plasma nitrite concentrations reported similar variability between study outcomes when assessing the disruption of the oral microbiota using antiseptic mouthwashes, with four of the six studies reporting a reduction of plasma nitrite concentrations, whilst for two studies there was no difference [39]. Furthermore, three studies out of five studies reported higher SBP in mouthwash groups compared to controls [39]. Following dietary nitrate supplementation, BP reductions have been correlated with increased plasma nitrite concentrations, implicating nitrate reduction as important in the efficacy of the intervention [38, 76]. Although there is a role for oral bacteria in *in vivo* nitrate cycling, the impact on plasma nitrite and NO, as well as subsequent cardiovascular responses, remains poorly understood.

A similar degree of unpredictability in responses to acute nitrate challenges was observed within the current study. Some oral microcosms displayed a clear increase in NaR activity and thus higher nitrite within the perfusate, alongside a shift in bacterial profiles towards high relative abundances of known nitrate reducers, whilst others did not respond. The results of this study reflect previous *in vivo* results regarding the unpredictability surrounding the efficacy of nitrate as a prebiotic in the treatment of cardiovascular disorders [39, 73]. In a previous *in vitro* investigation, which aimed to identify nitrate-reducing candidate probiotics, oral microcosms that did not initially respond to nitrate did respond once a high nitrate-reducing probiotic bacterium was introduced in tandem with the nitrate in short period (4-7 hours) experiments [77]. Preliminary identification of high NaR-producing bacterium with a strong safety profile, a minimal effective dose and the potential for large-scale industrial production with the possibility of incorporation in consumer dietary products should be studied *in vitro* and *in vivo* [78]. These studies should examine populations with both high and low baseline NaR activity to further investigate the preliminary evidence that those with lowest salivary NaR respond most to prebiotic nitrate that is presented in this current study. Furthermore, high nitrate-reducing oral bacterium has been suggested to have an anti-cariogenic effect, mediated by alkali production and lactate consumption limiting a drop in pH as carbohydrates are fermented [79-81]. Monitoring of medium pH *in vitro* would allow this hypothesis to be investigated further.

We have previously reported that the abundance and activity of oral nitrate-reducing bacteria impact influence blood pressure in hypertensive pregnant women and baseline blood pressure can be reduced by dietary nitrate supplementation [40]. Whilst the principle that oral nitrate metabolism can positively affect systemic health has been established, the mechanisms which underlie interindividual variation in responses are not fully understood and a system through which preclinical studies can be conducted of this and of possible interventions has several potential applications. The current study therefore aimed to establish and apply an *in vitro* oral model to assess the ability to manipulate community composition and associated nitrite formation following supplementation with inorganic nitrate. Whilst the nitrate concentration used to dose the models (15 mM) was broadly representative of physiological concentrations, it was relatively high [38]. The typical nitrate concentration range in the saliva is 5-8 mM typically after a nitrate-rich meal [85]. Secondly, fresh saliva as opposed to frozen saliva should be used as inoculum, although there was no change in alpha or beta diversity between the inoculum and the subsequent microcosms in the present study. Whilst losses in microbial diversity may occur when inoculum has been introduced into an *in vitro* model [15, 47], this was not observed in the current study.

*In vitro* modelling has previously facilitated the analysis of oral microcosms and ecological perturbations in numerous clinical disorders of the oral cavity, including caries [82] and periodontal disease [83] and subsequent anti-plaque interventions involving probiotics [84, 85], *L*-arginine [86, 87] and triclosan [88]. Continuous culture *in vitro* models involve the growth and maintenance of a consortium of bacteria following inoculation with representative poly-or mono-species microbial populations [89]. These systems enable continuous nutrient availability and the removal of detrimental metabolic waste products, consequently generating persistent and reproducible oral microcosms [43]. *In vitro* studies have been undertaken for the isolation of high nitrate-reducing bacterial species as possible probiotics (live organisms which, when consumed in sufficient quantities, confer a health benefit to the host) [90], alongside the assessment of nitrate itself as a prebiotic (a non-digestible food that can be used as a substrate by host microorganisms to produce energy, metabolites or micronutrients) [77, 80, 91].

The reduction of nitrate to nitrite in the oral cavity by host bacterium is associated with systemic health, but our current understanding of the underlying mechanisms is incomplete. Interindividual variability in the ability to reduce nitrate [73, 74] was reproduced in the microcosms. Such variability in this alternative entero-salivary pathway has implications in disorders where NO deficiency is implicated [92]. Greater appreciation of this symbiotic relationship and how pre- and probiotic inventions could shift the ecological balance to improve host health offers an opportunity in the treatment of cardiometabolic disorders. Our data suggest that prebiotic nitrate supplementation may be used to alter oral nitrate metabolism, which has been shown in previous *in vitro* [80] and *in vivo* studies [93], with dietary interventions an attractive therapeutic avenue for hypertension [94, 95]. Combination approaches involving high nitrate-reducing probiotic bacteria alongside prebiotic nitrate supplementation may be required in some individuals. A more complete understanding of the microbiology behind the metabolism of nitrate and nitrite in the oral cavity is now a priority [96].

## Author Contributions

T.W. collected the data. A.J.M., E.C., G.H., J.M. and T.W. designed the experiments. T.W., H.S. and P.C. analysed the data. T.W. and H.S. wrote the manuscript. All authors read and approved the final manuscript.

## Acknowledgements

The authors would like to thank the staff at the Maternal and Fetal Health Research Centre and the School of Pharmacy, University of Manchester.

## Data Availability Statement

The data that support the findings of this study are available from the corresponding author upon reasonable request. Sequence data will be openly available in the sequence read archive.

## References

1. NICE, Hypertension: the clinical management of primary hypertension in adults 2011 August (updated 2016 November):CG127. National Institute of Clinical Excellence. London (UK): National Clinical Guideline Centre, 2016.

2. Saiz, L.C., et al., Blood pressure targets for the treatment of people with hypertension and cardiovascular disease. The Cochrane database of systematic reviews, 2018. 7(7): p. CD010315–CD010315.

3. Forouzanfar, M.H., et al., Global Burden of Hypertension and Systolic Blood Pressure of at Least 110 to 115 mm Hg, 1990-2015. JAMA, 2017. 317(2): p. 165–182.

4. Dominiczak, A., C. Delles, and S. Padmanabhan, Genomics and Precision Medicine for Clinicians and Scientists in Hypertension. Hypertension, 2017. 69(4): p. e10–e13.

5. Lundberg, J.O., et al., Nitrate, bacteria and human health. Nature Reviews Microbiology, 2004. 2(7): p. 593–602.

6. Lundberg, J.O., E. Weitzberg, and M.T. Gladwin, The nitrate–nitrite–nitric oxide pathway in physiology and therapeutics. Nature Reviews Drug Discovery, 2008. 7: p. 156.

7. Grant, M.M. and D. Jönsson, Next Generation Sequencing Discoveries of the Nitrate-Responsive Oral Microbiome and Its Effect on Vascular Responses. J Clin Med, 2019. 8(8).

8. Alzahrani, H.S., et al., The role of dietary nitrate and the oral microbiome on blood pressure and vascular tone. Nutr Res Rev, 2021. 34(2): p. 222–239.

9. Remington, J. and K. Winters, Effectiveness of dietary inorganic nitrate for lowering blood pressure in hypertensive adults: a systematic review. JBI Database System Rev Implement Rep, 2019. 17(3): p. 365–389.

10. He, Y., et al., Effect of inorganic nitrate supplementation on blood pressure in older adults: A systematic review and meta-analysis. Nitric Oxide, 2021. 113-114: p. 13–22.

11. Hobbs, D.A., T.W. George, and J.A. Lovegrove, The effects of dietary nitrate on blood pressure and endothelial function: a review of human intervention studies. Nutrition Research Reviews, 2013. 26(2): p. 210–222.

12. Kalaycıoğlu, Z. and F.B. Erim, Nitrate and Nitrites in Foods: Worldwide Regional Distribution in View of Their Risks and Benefits. J Agric Food Chem, 2019. 67(26): p. 7205–7222.

13. Qin, L., et al., Sialin functions as a nitrate transporter in the plasma membrane. Proceedings of the National Academy of Sciences, 2012. 109(33): p. 13434.

14. Qu, X.M., et al., From Nitrate to Nitric Oxide: The Role of Salivary Glands and Oral Bacteria. Journal of Dental Research, 2016. 95(13): p. 1452–1456.

15. Doel, J.J., et al., Evaluation of bacterial nitrate reduction in the human oral cavity. 2005. 113(1): p. 14–19.

16. Ahmed, K.A., et al., Measuring nitrate reductase activity from human and rodent tongues. Nitric Oxide, 2017. 66: p. 62–70.

17. Panza, J., et al., Panza JA, Quyyumi AA, Brush Jr JE, Epstein SE. Abnormal endothelium-dependent vascular relaxation in patients with essential hypertension. N Engl J Med 323, 22-27. Vol. 323. 1990. 22–7.

18. Chowienczyk, P.J., et al., Impaired endothelium-dependent vasodilation of forearm resistance vessels in hypercholesterolaemia. The Lancet, 1992. 340(8833): p. 1430–1432.

19. Bhupathiraju, S.N., et al., Quantity and variety in fruit and vegetable intake and risk of coronary heart disease. Am J Clin Nutr, 2013. 98(6): p. 1514–23.

20. Ahmad, A., et al., Role of Nitric Oxide in the Cardiovascular and Renal Systems. Int J Mol Sci, 2018. 19(9).

21. Hsu, C.N. and Y.L. Tain, Regulation of Nitric Oxide Production in the Developmental Programming of Hypertension and Kidney Disease. Int J Mol Sci, 2019. 20(3).

22. Alzahrani, H.S., et al., The role of dietary nitrate and the oral microbiome on blood pressure and vascular tone. Nutr Res Rev, 2020: p. 1–18.

23. Koch, C.D., et al., Enterosalivary nitrate metabolism and the microbiome: Intersection of microbial metabolism, nitric oxide and diet in cardiac and pulmonary vascular health. Free Radical Biology and Medicine, 2017. 105: p. 48–67.

24. Kapil, V., et al., Physiological role for nitrate-reducing oral bacteria in blood pressure control. Free radical biology & medicine, 2013. 55: p. 93–100.

25. McDonagh, S.T., et al., The Effects of Chronic Nitrate Supplementation and the Use of Strong and Weak Antibacterial Agents on Plasma Nitrite Concentration and Exercise Blood Pressure. Int J Sports Med, 2015. 36(14): p. 1177–85.

26. Sundqvist, M.L., J.O. Lundberg, and E. Weitzberg, Effects of antiseptic mouthwash on resting metabolic rate: A randomized, double-blind, crossover study. Nitric Oxide, 2016. 61: p. 38–44.

27. Woessner, M., et al., A stepwise reduction in plasma and salivary nitrite with increasing strengths of mouthwash following a dietary nitrate load. Nitric Oxide, 2016. 54: p. 1–7.

28. Senkus, K.E. and K.M. Crowe-White, Influence of mouth rinse use on the enterosalivary pathway and blood pressure regulation: A systematic review. Crit Rev Food Sci Nutr, 2020. 60(17): p. 2874–2886.

29. Bryan, N.S., G. Tribble, and N.J.C.H.R. Angelov, Oral Microbiome and Nitric Oxide: the Missing Link in the Management of Blood Pressure. 2017. 19(4): p. 33.

30. Webb, A.J., et al., Acute blood pressure lowering, vasoprotective, and antiplatelet properties of dietary nitrate via bioconversion to nitrite. Hypertension (Dallas, Tex. : 1979), 2008. 51(3): p. 784–790.

31. Hobbs, D.A., et al., Blood pressure-lowering effects of beetroot juice and novel beetrootenriched bread products in normotensive male subjects. British Journal of Nutrition, 2012. 108(11): p. 2066–2074.

32. Kapil, V., et al., Dietary nitrate provides sustained blood pressure lowering in hypertensive patients: a randomized, phase 2, double-blind, placebo-controlled study. Hypertension (Dallas, Tex. : 1979), 2015. 65(2): p. 320–327.

33. Velmurugan, S., et al., Dietary nitrate improves vascular function in patients with hypercholesterolemia: a randomized, double-blind, placebo-controlled study. The American journal of clinical nutrition, 2016. 103(1): p. 25–38.

34. Avoort, C., et al., Increasing Nitrate-Rich Vegetable Intake Lowers Ambulatory Blood Pressure in (pre)Hypertensive Middle-Aged and Older Adults: A 12-Wk Randomized Controlled Trial. J Nutr, 2021.

35. Gilchrist, M., et al., Effect of dietary nitrate on blood pressure, endothelial function, and insulin sensitivity in type 2 diabetes. Free Radical Biology and Medicine, 2013. 60: p. 89–97.

36. Bondonno, C.P., et al., Absence of an effect of high nitrate intake from beetroot juice on blood pressure in treated hypertensive individuals: a randomized controlled trial. The American Journal of Clinical Nutrition, 2015. 102(2): p. 368–375.

37. Burleigh, M.C., et al., Salivary nitrite production is elevated in individuals with a higher abundance of oral nitrate-reducing bacteria. Free Radical Biology and Medicine, 2018. 120: p. 80–88.

38. Ormesher, L., et al., Effects of dietary nitrate supplementation, from beetroot juice, on blood pressure in hypertensive pregnant women: A randomised, double-blind, placebo-controlled feasibility trial. Nitric Oxide, 2018. 80: p. 37–44.

39. Zhurakivska, K., et al., Do Changes in Oral Microbiota Correlate With Plasma Nitrite Response? A Systematic Review. Frontiers in Physiology, 2019. 10(1029).

40. Willmott, T., et al., Altered Oral Nitrate Reduction and Bacterial Profiles in Hypertensive Women Predict Blood Pressure Lowering Following Acute Dietary Nitrate Supplementation. Hypertension, 2023.

41. Greenman, J., et al., In vitro models for oral malodor. Oral Dis, 2005. 11 Suppl 1: p. 14–23.

42. McBain, A.J., et al., Development and characterization of a simple perfused oral microcosm. J Appl Microbiol, 2005. 98(3): p. 624–34.

43. McBain, A.J., Chapter 4: In vitro biofilm models: an overview. Adv Appl Microbiol, 2009. 69: p. 99–132.

44. Ledder, R.G., et al., An in vitro evaluation of hydrolytic enzymes as dental plaque control agents. J Med Microbiol, 2009. 58(Pt 4): p. 482–491.

45. Ledder, R.G. and A.J. McBain, An in vitro comparison of dentifrice formulations in three distinct oral microbiotas. Arch Oral Biol, 2012. 57(2): p. 139–47.

46. Ledder, R.G., et al., Individual microflora beget unique oral microcosms. J Appl Microbiol, 2006. 100(5): p. 1123–31.

47. Koopman, J.E., et al., Nitrate and the Origin of Saliva Influence Composition and Short Chain Fatty Acid Production of Oral Microcosms. Microbial ecology, 2016. 72(2): p. 479–492.

48. Lowe, R.H. and H.J. Evans, Preparation and some properties of a soluble nitrate reductase from Rhizobium japonicum. Biochimica et Biophysica Acta (BBA) - Specialized Section on Enzymological Subjects, 1964. 85(3): p. 377–389.

49. Allison, C. and G.T. Macfarlane, Dissimilatory nitrate reduction by Propionibacterium acnes. Applied and environmental microbiology, 1989. 55(11): p. 2899–2903.

50. Caporaso, J.G., et al., Ultra-high-throughput microbial community analysis on the Illumina HiSeq and MiSeq platforms. The ISME journal, 2012. 6(8): p. 1621–1624.

51. Rullo, J., et al., Local oral and nasal microbiome diversity in age-related macular degeneration. Scientific Reports, 2020. 10(1): p. 3862.

52. Demmitt, B.A., et al., Genetic influences on the human oral microbiome. BMC Genomics, 2017. 18(1): p. 659.

53. Granato, D.C., et al., Meta-omics analysis indicates the saliva microbiome and its proteins associated with the prognosis of oral cancer patients. Biochimica et Biophysica Acta (BBA) - Proteins and Proteomics, 2021. 1869(8): p. 140659.

54. Su, S.-C., et al., Oral microbial dysbiosis and its performance in predicting oral cancer. Carcinogenesis, 2020. 42(1): p. 127–135.

55. Caporaso, J.G., et al., QIIME allows analysis of high-throughput community sequencing data. Nature methods, 2010. 7(5): p. 335–336.

56. Bolyen, E., et al., Reproducible, interactive, scalable and extensible microbiome data science using QIIME 2. Nat Biotechnol, 2019. 37(8): p. 852–857.

57. Callahan, B.J., et al., Bioconductor Workflow for Microbiome Data Analysis: from raw reads to community analyses. F1000Research, 2016. 5: p. 1492–1492.

58. Bokulich, N.A., et al., Optimizing taxonomic classification of marker-gene amplicon sequences with QIIME 2’s q2-feature-classifier plugin. Microbiome, 2018. 6(1): p. 90.

59. McDonald, D., et al., An improved Greengenes taxonomy with explicit ranks for ecological and evolutionary analyses of bacteria and archaea. The ISME Journal, 2012. 6(3): p. 610–618.

60. Bisanz, J.E., qiime2R: Importing QIIME2 artifacts and associated data into R sessions. 2018. 0.99.

61. McMurdie, P.J. and S. Holmes, Phyloseq: a bioconductor package for handling and analysis of high-throughput phylogenetic sequence data. Pacific Symposium on Biocomputing. Pacific Symposium on Biocomputing, 2012: p. 235–246.

62. Wickham, H., ggplot2: Elegant Graphics for Data Analysis. Springer-Verlag New York, 2016.

63. Team, R.C., R: A Language and Environment for Statistical Computing. R Foundation for Statistical Computing. 2016.

64. Love, M.I., W. Huber, and S. Anders, Moderated estimation of fold change and dispersion for RNA-seq data with DESeq2. Genome Biology, 2014. 15(12): p. 550.

65. Kolenbrander, P.E., et al., Oral multispecies biofilm development and the key role of cell-cell distance. Nat Rev Microbiol, 2010. 8(7): p. 471–80.

66. Chen, H., et al., Microbial characteristics across different tongue coating types in a healthy population. Journal of Oral Microbiology, 2021. 13(1): p. 1946316.

67. Rosier, B.T., et al., Nitrate as a potential prebiotic for the oral microbiome. Scientific Reports, 2020. 10(1): p. 12895.

68. Rosier, B.T., et al., Isolation and Characterization of Nitrate-Reducing Bacteria as Potential Probiotics for Oral and Systemic Health. Frontiers in Microbiology, 2020. 11.

69. Engel, A.S., et al., Biofilm formation on different dental restorative materials in the oral cavity. BMC Oral Health, 2020. 20(1): p. 162.

70. Kuboniwa, M. and R.J. Lamont, Subgingival biofilm formation. Periodontol 2000, 2010. 52(1): p. 38–52.

71. Liddle, L., et al., Variability in nitrate-reducing oral bacteria and nitric oxide metabolites in biological fluids following dietary nitrate administration: An assessment of the critical difference. Nitric Oxide, 2019. 83: p. 1–10.

72. Sato-Suzuki, Y., et al., Nitrite-producing oral microbiome in adults and children. Scientific Reports, 2020. 10(1): p. 16652.

73. Blekkenhorst, L.C., et al., Nitrate, the oral microbiome, and cardiovascular health: a systematic literature review of human and animal studies. Am J Clin Nutr, 2018. 107(4): p. 504–522.

74. Li, D., et al., Repeated administration of inorganic nitrate on blood pressure and arterial stiffness: a systematic review and meta-analysis of randomized controlled trials. Journal of hypertension, 2020. 38(11): p. 2122–2140.

75. Remington, J. and K. Winters, Effectiveness of dietary inorganic nitrate for lowering blood pressure in hypertensive adults: a systematic review. JBI Evidence Synthesis, 2019. 17(3): p. 365–389.

76. Kapil, V., et al., Inorganic nitrate supplementation lowers blood pressure in humans: role for nitrite-derived NO. Hypertension, 2010. 56(2): p. 274–81.

77. Rosier, B.T., et al., Isolation and Characterization of Nitrate-Reducing Bacteria as Potential Probiotics for Oral and Systemic Health. Frontiers in microbiology, 2020. 11: p. 555465–555465.

78. Fenster, K., et al., The Production and Delivery of Probiotics: A Review of a Practical Approach. Microorganisms, 2019. 7(3).

79. Li, H., et al., Salivary nitrate--an ecological factor in reducing oral acidity. Oral Microbiol Immunol, 2007. 22(1): p. 67–71.

80. Rosier, B.T., et al., Nitrate as a potential prebiotic for the oral microbiome. Scientific reports, 2020. 10(1): p. 12895–12895.

81. Rosier, B.T., et al., A Single Dose of Nitrate Increases Resilience Against Acidification Derived From Sugar Fermentation by the Oral Microbiome. Frontiers in cellular and infection microbiology, 2021. 11: p. 692883–692883.

82. Wu, J.-S., et al., Porphyromonas gingivalis Promotes 4-Nitroquinoline-1-Oxide-Induced Oral Carcinogenesis With an Alteration of Fatty Acid Metabolism. Frontiers in microbiology, 2018. 9: p. 2081–2081.

83. Velsko, I.M. and L.M. Shaddox, Consistent and reproducible long-term in vitro growth of health and disease-associated oral subgingival biofilms. BMC microbiology, 2018. 18(1): p. 70–70.

84. Schwendicke, F., et al., Inhibition of Streptococcus mutans Growth and Biofilm Formation by Probiotics in vitro. Caries Res, 2017. 51(2): p. 87–95.

85. Humphreys, G.J. and A.J. McBain, Antagonistic effects of Streptococcus and Lactobacillus probiotics in pharyngeal biofilms. Lett Appl Microbiol, 2019. 68(4): p. 303–312.

86. Kolderman, E., et al., L-arginine destabilizes oral multi-species biofilm communities developed in human saliva. PLoS One, 2015. 10(5): p. e0121835.

87. Ledder, R.G., et al., Arginine Exposure Decreases Acidogenesis in Long-Term Oral Biofilm Microcosms. mSphere, 2017. 2(4): p. e00295–17.

88. Fernández, E., et al., Antibacterial Effects of Toothpastes Evaluated in an In Vitro Biofilm Model. Oral Health Prev Dent, 2017. 15(3): p. 251–257.

89. Brown, J.L., et al., Polymicrobial oral biofilm models: simplifying the complex. Journal of Medical Microbiology, 2019. 68(11): p. 1573–1584.

90. Rijkers, G.T., et al., Guidance for substantiating the evidence for beneficial effects of probiotics: current status and recommendations for future research. J Nutr, 2010. 140(3): p. 671s–6s.

91. Corzo, N., et al., Prebiotics: concept, properties and beneficial effects. Nutr Hosp, 2015. 31 Suppl 1: p. 99–118.

92. Lundberg, J.O., M. Carlström, and E. Weitzberg, Metabolic Effects of Dietary Nitrate in Health and Disease. Cell Metabolism, 2018. 28(1): p. 9–22.

93. Vanhatalo, A., et al., Nitrate-responsive oral microbiome modulates nitric oxide homeostasis and blood pressure in humans. Free radical biology & medicine, 2018. 124: p. 21–30.

94. Ashor, A., J. Lara, and M. Siervo, Medium-term effects of dietary nitrate supplementation on systolic and diastolic blood pressure in adults: A systematic review and meta-analysis. Vol. 35. 2017. 1.

95. O’Gallagher, K., et al., Grapefruit juice enhances the systolic blood pressure-lowering effects of dietary nitrate-containing beetroot juice. British Journal of Clinical Pharmacology, 2021. 87(2): p. 577–587.

96. Goh, C.E., et al., Association Between Nitrate-Reducing Oral Bacteria and Cardiometabolic Outcomes: Results From ORIGINS. J Am Heart Assoc, 2019. 8(23): p. e013324.

